## Supplemental Information for "Alternatives to Friction Coefficient: Fine Touch Perception Correlates with Frictional Instabilities"

**Supporting Information for**
### Alternatives to Friction Coefficient: Role of Frictional Instabilities on Fine Touch Perception

Maryanne Derkaloustian[a], Pushpita Bhattacharyya[b], Truc Ngo[c], Joshua G. A. Cashaback[c], Jared Medina[b,d], Charles B. Dhong[a,c*]

[a]*Department of Materials Science and Engineering, University of Delaware, Newark, DE, USA*
[b]*Department of Psychological and Brain Sciences, University of Delaware, Newark, DE, USA*
[c]*Department of Biomedical Engineering, University of Delaware, Newark, DE, USA*
[d]*Department of Psychology, Emory University, Atlanta, GA, USA*

#### Atomic Force Microscopy

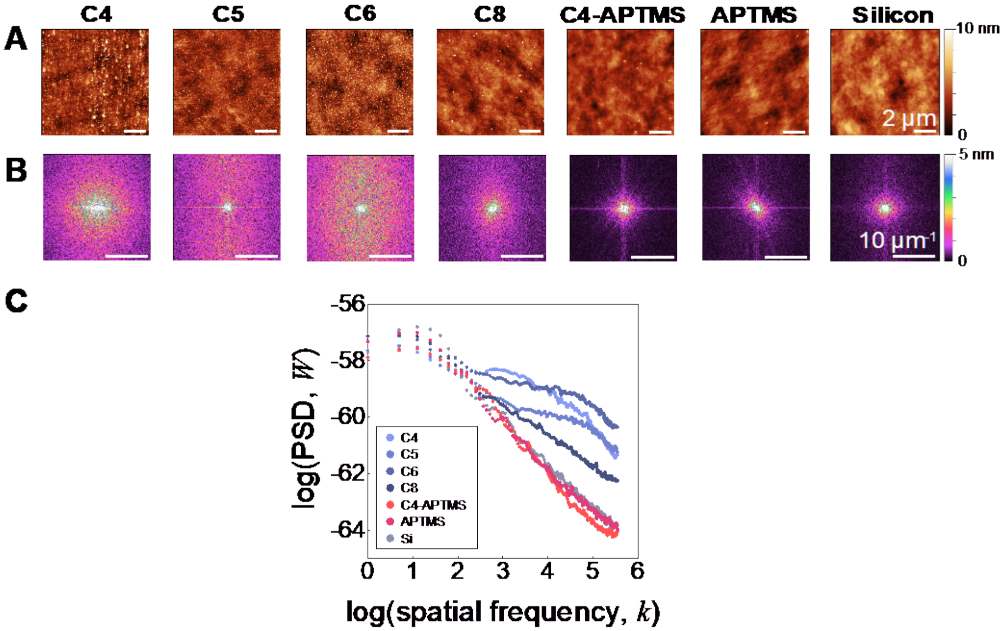


**Fig. S1.** Surface characterization via atomic force microscopy. A) Height profiles of surfaces obtained via AFM in tapping mode. All scale bars are 2 µm. B) Respective 2D fast Fourier transforms of surface topographies. All scale bars are 10 µm$^{-1}$. C) 2D power spectral densities of all surface profiles, plotted as log(PSD) vs. log(spatial frequency) to calculate Hurst exponents.

Average roughness ($R_a$) of the AFM topographies in **Fig. S1A** were calculated along their diagonals (~14.14 µm). The values in **Table 1** are all on the small length scale of 100 pm, although **C4** and **C6** are slightly higher end due to the formation of more aggregates. This trend does not hold with **C8**, however, as the longer chains more easily form ordered films with less drastic height changes overall.[1] All alkylsilanes form a rougher coating than the aminosilanes, also consistent with literature.[2]

The two-dimensional fast Fourier transforms (2D FFTs) of the height profiles shown in **Fig. S1B** are largely uniform. The shorter chain alkylsilanes present more diffuse rings in Fourier space, indicative of a lack of self-similarity across length scales.[3] Ordering and more fractal behavior begin to emerge with **C8**, as expected of longer chains with increased intermolecular interactions.[1,2] Meanwhile, the aminosilanes most closely replicate the underlying Si substrate in their isotropic, fractal behavior.

2D power spectral densities (PSDs) of the silanes were plotted in **Fig. S1C** to quantify their scaling behavior through the Hurst exponent, *H*. This exponent identifies how roughness evolves across length scales: a lower *H* between 0 and 0.5 corresponds to a surface with a more homogeneous distribution of continuous high and low features, while *H* between 0.5 and 1 indicates the changes in roughness are sharper.[2,4] This is calculated as $H = 0.5 \times (|\text{slope}| - 1)$, using the slope of each surface's linear regime. A broad linear regime of smaller slope is observed for bare Si, as well as **C8**, **C4-APTMS**, and **APTMS**, resulting in lower *H* (see **Table 1**). This is consistent with previous works, as van der Waals forces from increased chain length and hydrogen bonding from the amine groups both allow for chain alignment, with the roughness profile of uncoated Si well maintained and ordering preserved across most length scales.[2] This is in contrast to the profiles observed for **C4**, **C5**, and **C6**: multiple regimes appear, with steeper slopes observed at higher spatial frequencies. **C5** and **C6** still exhibit $H < 0.5$, but are self-affine to a lower extent than **C8** and the aminosilane surfaces. **C4** demonstrates more abrupt changes in height, even excluding the effects of the larger polymer aggregates ($H = 0.55$ with masking). These higher *H* thus represent increased disorder, including gauche defects and horizontal chain alignment,[5] masking the underlying topography of bare Si.

**X-ray Photoelectric Spectroscopy**

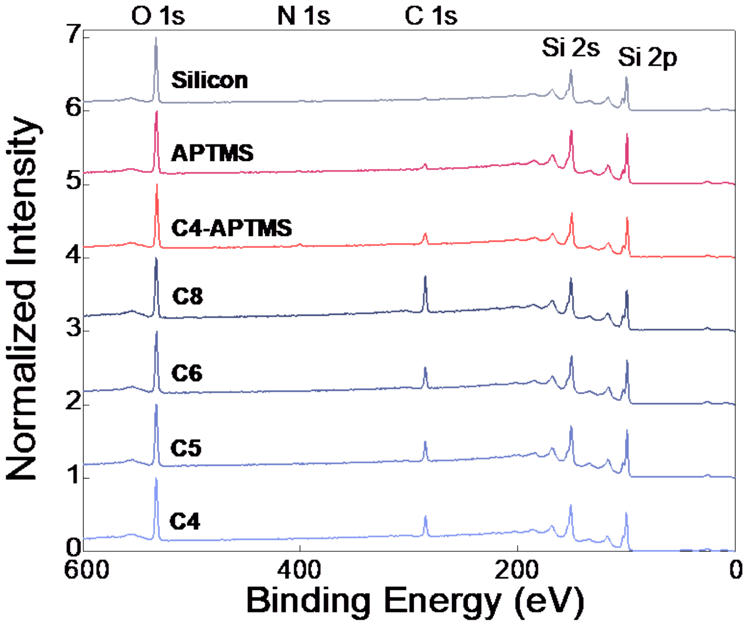


**Fig. S2.** XPS survey scans of surfaces. C, N, and O peaks indicate evidence of binding onto Si wafers after deposition. Plotted as intensities normalized by the O peaks vs. binding energy.

XPS survey scans in **Fig. S2** identify the elements present on our surfaces with a quantifiable photoelectric signal. To compare surfaces, all spectra were normalized by the highest peak with no expected change, corresponding to oxygen with a 532 eV peak.[6] Silanization is evident through the appearance of a carbon (285 eV) peak,[7] of heights incrementally increasing with chain length in the alkylsilane surfaces (**C4**: 0.48, **C5**: 0.51, **C6**: 0.52, and **C8**: 0.75 × the height of the O peak). The C peaks are smaller but still visible in aminosilanes (**C4-APTMS**: 0.33 and **APTMS**: 0.30 × O). Small nitrogen peaks at 400 eV (0.19 × O) also appear in these two surfaces.[8] Most importantly, the spectra present evidence of covalent bonding to the surfaces and successful deposition instead of physisorption. In the unreacted state, the alkylsilanes are chlorine-terminated, but no chlorine peak at 200 eV is present in the final surfaces.[7,9] Unreacted aminosilanes end with a methoxy group, but the C peaks from their deposited state are still small.

**Water Contact Angle Hysteresis**

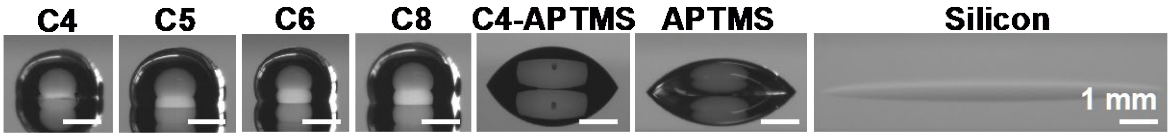


**Fig. S3.** DI water droplets on surfaces, representative images used to measure advancing angles. All scale bars are 1 mm.

Even with differences in the AFM results, all the surfaces exhibit low chemical heterogeneity,[2,10] with water contact angle hysteresis < 10º. There are noticeable differences in hydrophobicity, shown in the images used to measure advancing angles in **Fig. S3**. The alkylsilanes form considerably rounder droplets indicating a higher degree of hydrophobicity than the aminosilanes.[11] However, even the aminosilanes are still more hydrophobic than the plasma-treated, uncoated Si, indicating the availability of free O allowing silanes to react and bind to the surfaces when undergoing depositions.[12]

**Instability Classification**

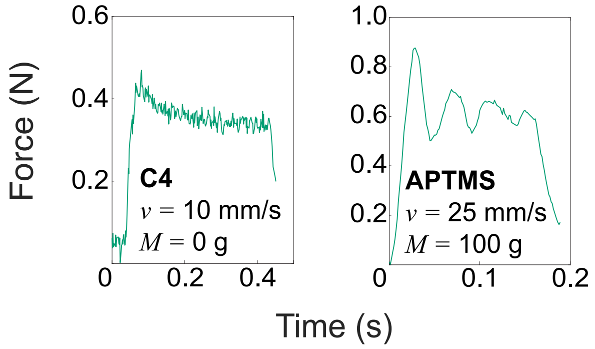


**Fig. S4**. Force traces with less distinct categorization. Left: stiction spike with small height, right: stiction spike followed by small oscillations.

In many cases, instability classification was straightforward. However, in cases where stiction spikes where not as prominent as those in **Fig. 1B**, only those at least 10% above the mean subsequent steady sliding were classified as stiction spikes. However, when the remainder of the trace still oscillates, a stiction spike was counted only if it was 40% higher in magnitude than the mean of the traces. Examples of these visually categorized stiction spikes are shown in **Fig. S4**. Most importantly, these in-between cases mostly exist at boundary zones on our phase maps, where the same mass-velocity combinations do not produce identical force profiles. The boundaries do not only represent sharp transitions between instability phases, but also "mixed-case" force traces.

**Statistical Model Selection**

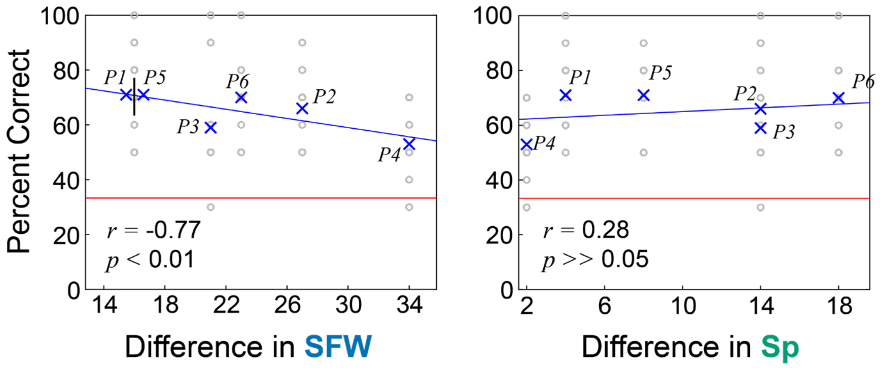


**Fig. S5.** GLMM fits of participant accuracy vs. the differences in instability incidence for individual instability types. Left: accuracy vs. differences in formation of slow frictional waves (SFW) between pairs. *P1* and *P5* have the same *x*-axis value and are shifted for clarity. Right: accuracy vs. differences in formation of stiction spikes (Sp).

Participant data was fit to generalized linear mixed models (GLMMs) in order to decouple the variability of fixed and random effects, and for overall higher statistical power.[13] We first assessed subject accuracy by fitting normally distributed accuracy to differences in each instability type between pairs, as we hypothesized that all three instability types were important in predicting human performance. When examining each type individually, we first observed a positive, statistically significant relationship between accuracy and steady sliding ($p < 0.05$). We also saw a negative correlation between accuracy and slow frictional waves that was statistically significant ($p < 0.01$), and no correlation between accuracy and stiction spikes (**Fig. S5**). We attribute these findings to the locations of these instability zones on the material phase maps. However, fitting all instability differences as terms in one GLMM did not yield any statistically significant results. A similar approach was used to correlate response times to instability types, which showed a statistically significant, negative correlation between response time and stiction spikes (details in **main text**).

**Effects of Material Properties**

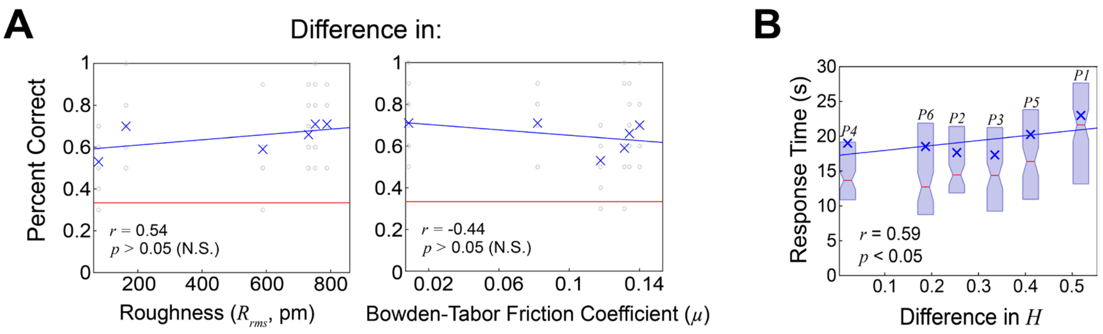


**Fig. S6.** GLMMs of accuracy vs. more material parameters or properties across N = 10 participants. A) Left: root-mean-square roughness $R_{rms}$, right: velocity-dependent Bowden-Tabor friction coefficient across all conditions ($\mu$). B) GLMM fit of response times vs. differences in Hurst exponent $H$. Mean times are represented by blue × marks, while medians are represented by red lines at notches of box plots.

After determining that $R_a$ was not a strong predictor, we also performed a GLMM fit on root-mean-square roughness $R_{rms}$, to verify that any common definition of roughness does not explain human performance. Although $R^2$ was slightly improved for this measure of roughness, each pair was located similarly to $R_a$, leading to a fit that was still not statistically significant (**Fig. S6A**).

The Bowden-Tabor friction coefficient of each material was determined by plotting average friction force vs. normal force for each velocity. Normal force was approximated as the applied load + the deadweight of the mock finger (6 g). Based on the equation $F = \mu N + \sigma A$, the slope of the points determined velocity-dependent friction coefficients,[14] which were then averaged to obtain a singular average friction coefficient for each material. Interestingly, these led to a negative fit similar to the more simplified friction coefficient, but is not statistically significant.

As a positive correlation between accuracy and Hurst exponent was observed ($p < 0.05$, details in main text), we fit the mean response times of each pair similarly. Here, we saw a positive, statistically significant relationship again ($p < 0.05$, **Fig. S6B**) , meaning the differences between pairs slow participants down. Unlike stiction spikes and frictional instabilities more broadly, each material's Hurst exponent is static, and participants cannot modulate their exploration conditions to feel other $H$ on the same material. No transitions exist to trigger a response, although after increased time participants do successfully distinguish between $H$ on the silane surfaces.
